# Assembly of hundreds of novel bacterial genomes from the chicken caecum

**DOI:** 10.1101/699843

**Authors:** Laura Glendinning, Robert D. Stewart, Mark J. Pallen, Kellie A. Watson, Mick Watson

## Abstract

Chickens are a highly important source of protein for a large proportion of the human population. The caecal microbiota plays a crucial role in chicken nutrition through the production of short chain fatty acids, nitrogen recycling and amino acid production. In this study we sequenced DNA from caecal contents samples taken from 24 chickens belonging to either a fast or slower growing breed consuming either a vegetable-only diet or a diet containing fish meal. We utilised 1.6T of Illumina data to construct 469 draft metagenome-assembled bacterial genomes, including 460 novel strains, 283 novel species and 42 novel genera. We compared our genomes to data from eight EU countries and show that these genomes are abundant within European chicken flocks. We also compared the abundance of our genomes, and the carbohydrate active enzymes they produce, between our chicken groups and demonstrate that there are both breed- and diet-specific microbiomes, as well as an overlapping core microbiome. This data will form the basis for future studies examining the composition and function of the chicken caecal microbiota.

## Background

There are an estimated 23 billion live chickens on the planet at any one time (1), out-numbering humans by over 3:1. As most of these are reared for food, the actual number of chickens produced per year is even higher, at almost 65 billion, leading some to speculate that the accumulation of chicken bones in the fossil record will be used by future archaeologists as a unique marker for the Anthropocene (2).

Since the 1960s, worldwide chicken meat production has increased by over ten times (3). Global meat production is predicted to be 16% higher in 2025 vs. 2015, with most of this increase originating from poultry meat production (4). Part of the popularity of chicken meat is that, due to intensive selection, chickens have been developed which are highly productive in terms of their growth rate with efficient feed conversion ratios (the rate at which chickens convert feed into muscle), decreasing from 3.0 in the 1960s to 1.7 in 2005 (5), making them a cheap source of protein in comparison to other livestock. Chickens also produce less greenhouse gasses per kg of meat than pigs, cattle and sheep (6). The potential to manipulate the microbiota in chickens to gain further increases in productivity is of great commercial and scientific interest, leading to the use of probiotics across the poultry industry (7).

As well as playing an important role in pathogen protection (8) and immune system development (9), the microbiota of the chicken also plays a crucial nutritional role. The largest concentration of microbial cells in the chicken gastrointestinal tract can be found in the caeca and thereby the majority of chicken microbiota studies focus primarily on these microbial communities. Members of the caecal microbiota are able to produce short chain fatty acids (SCFAs) such as acetate, butyrate, lactate and propionate from carbohydrate sources which have passed through the small intestine; these SCFAs are then able to be absorbed by the bird and used as an energy source (10). Members of the chicken caecal microbiota have also been implicated in the recycling of nitrogen by the degradation of nitrogenous compounds (11) and the synthesis of amino acids (12). One study demonstrated that 21% of the variation in chicken abdominal fat mass could be attributed to the caecal microbiota composition, when controlling for host genetic effects (13). Differences have also been observed between birds with high and low feed efficiency (14, 15). However, despite extensive research over many decades, the quantitative importance of the caeca in chicken nutrition remains unclear (16), and relatively few microbes commensal in the chicken gut have been sequenced and deposited in public repositories.

The emergence of cheaper DNA sequencing technologies (17, 18) has led to an explosion in studies which have sought to characterise the chicken gastrointestinal microbiota, particularly using 16S rRNA gene based methods. Using this methodology, it has been found that the chicken caecal microbiota in the first few weeks of life is predominantly colonised by members of the *Firmicutes*, mostly of the order *Clostridiales* (8, 19). Whilst valuable, marker-gene studies do not enable an in-depth functional and genomic characterisation of the microbiome. Some microbes from the chicken caeca have been successfully cultured and sequenced, including 133 gut anaerobe strains representing a few dozen species with a wide range of metabolic potentials (20); however it is highly unlikely that these microbes represent the entire diversity of the chicken caecal microbiota, due to the difficulty in culturing many anaerobic gut microorganisms. One method which avoids this issue of culturability is the construction of metagenome assembled genomes (MAGs). Due to improvements in computational power and sequencing technologies, and the development of new computational approaches (21, 22), it is now possible to accurately bin short-read metagenomic data into high-quality genomes. Using this technique thousands of MAGs have been generated from various environments, including humans (23, 24), chickens (25) the rumen (26, 27), pig faeces (28), marine surface waters (29, 30), an underground aquifer system (31) and other public datasets (32).

In this study we sought to use metagenomic sequencing, assembly and binning to investigate the chicken caecal microbiota. In order to maximise diversity, we chose two commercial bird genotypes with different growth phenotypes, fed two different diets. This also allows us to look at the effects of breed and diet on strain level microbial abundance. The lines chosen for the study are Ross 308, a fast growing broiler breed, and the Ranger Classic, a slower growing broiler aimed at free-range, organic farms. All birds were fed either a vegetable-only diet or a diet based on fish meal as the protein source. The inclusion of fish meal in chicken diets has previously been linked to changes in the caecal microbiota and is correlated with an increased risk of necrotic enteritis (33, 34). We assemble 460 novel microbial strains, predicted to represent 283 novel microbial species and 42 novel microbial genera from the chicken microbiome, and go on to demonstrate both a breed- and diet-specific microbiota. We also demonstrate that our microbial genomes are abundant within European chicken flocks and represent the majority of reads from eight farms which were part of a pan-EU study examining antimicrobial resistance (AMR) in broilers (35). Whilst we show that large numbers of strains are shared between our birds, it is their relative abundance that largely drives breed and diet effects. This is the first large-scale binning of the chicken caecal microbiota, and we believe these data will form the basis for future studies of the structure and function of the chicken gut microbiome.

## Methods

### Ethical statement

Animals were housed in premises licensed under a UK Home Office Establishment License within the terms of the UK Home Office Animals (Scientific Procedures) Act 1986. Housing and husbandry complied with the Code of Practice for Housing and Care of Animals Bred, Supplied or Used for Scientific Purposes and were overseen by the Roslin Institute Animal Welfare and Ethical Review Board. Animals were culled by schedule one methods authorized by the Animals (Scientific Procedures) Act 1986.

### Study design

Ross 308 (Aviagen, UK) (n=12) and Ranger Classic (Aviagen, UK) (n=12) chickens were hatched and housed at the National Avian Research Facility in Edinburgh (UK). Birds were fed either a vegetable only diet or a diet supplemented with fish meal (**Table 1, Supplementary table 1**) (Diet formulation: **Supplementary tables 2 and 3**, nutritional info: **Supplementary table 4**). Birds received Mareks-Rispins vaccinations (Merial, France) at 1-2 days of age and were housed by group in separate floor pens (within the same room) with wood shaving bedding, and receiving food and water ad libitum. Stocking densities were based on UK Home Office Animals (Scientific Procedures) Act 1986, resulting in a floor area per bird of 0.133 m^2^ at 5 weeks of age. Birds were euthanized by cervical dislocation at 5 weeks of age and caecal content samples were collected. Contents from both caeca were pooled to make one sample per bird. Samples were stored at 4°C for a maximum of 24 hours until DNA extraction, except for those from DNA extraction batch 2 which were frozen at −20°C for 9 days prior to DNA extraction (**Supplementary table 5**). DNA extraction was performed as described previously using the DNeasy PowerLyzer PowerSoil Kit (Qiagen, UK) (36). Shotgun sequencing was performed on a NovaSeq (Illumina) producing 150bp paired-end reads.

**Table 1:** Chicken details

| Line | Diet | N | Mean body weight (kg) |
| --- | --- | --- | --- |
| Ranger Classic | Fish meal diet | 6 (3 male, 3 female) | Female: 1.89<br>Male: 2.13 |
|  | Vegetable diet | 6 (3 male, 3 female) | Female: 1.57<br>Male: 1.94 |
| Ross 308 | Fish meal diet | 6 (3 male, 3 female) | Female: 2.33<br>Male: 2.70 |
|  | Vegetable diet | 6 (4 male, 2 female) | Female: 2.23<br>Male: 2.89 |

### Bioinformatics

Assembly and binning was carried out as previously described (26, 27). Illumina adaptors were removed using trimmomatic (37). Single sample assemblies were performed using IDBA-UD (38) with the options --num_threads 16 --pre_correction --min_contig 300. BWA MEM (39) was used to separately map reads from every sample back to every assembly. SAMtools (40) was used to create BAM files and the command jgi_summarize_bam_contig_depths was run on all BAM files for each assembly to calculate coverage. A coassembly was also carried out on all 24 samples using MEGAHIT (options: --continue --kmin-1pass -m 100e+10 --k-list 27,37,47,57,67,77,87 --min-contig-len 1000 -t 16) (41). Contigs were filtered to a minimum length of 2kb, then indexed and mapped as for single assemblies.

METABAT2 (22) was used on both single-sample assemblies and co-assemblies to carry out metagenomic binning, taking into account coverage values and with the options --minContigLength 2000, --minContigDepth 2. All bins were dereplicated using dRep (42) with the options dereplicate_wf -p 16 -comp 80 -con 10 -str 100 -strW. Bins were dereplicated at 99% average nucleotide identity (ANI), resulting in each MAG being taxonomically equivalent to a microbial strain. Bins were also dereplicated at 95% ANI to calculate the number of species represented within our MAGs. CompareM was used to calculate average amino acid identity (AAI) (43).

The completeness and contamination of all bins was assessed using CheckM (44) with the options lineage_wf, -t 16, -x fa and filtering for completeness ≥80% and contamination ≤10%. GTDB-Tk (45) was used to assign taxonomy to MAGs, except for CMAG_333 which upon visual inspection of taxonomic trees was identified more accurately as *Clostridia*. For submission of our MAGs to NCBI, MAGs were named based on the following rule: if the lowest taxonomy assigned by GTDB-Tk did not correlate with an NCBI classification at the correct taxonomic level then MAGs were named after the lowest taxonomic level at which NCBI and GTDB-Tk matched. Comparative genomics between the MAGs and public datasets was carried out using MAGpy (46). The taxonomic tree produced by MAGpy was re-rooted manually using Figtree (47) at the branch between Firmicutes and the other bacterial phyla, and subsequently visualised using Graphlan (48). The novelty of genomes in comparison to those present in public databases was also determined. Genomes were defined as novel strains if the ANI output by GTDB-Tk was <99%. Genomes were determined as novel species if the ANI output by GTDB-Tk was <95% or if an ANI was not output by GTDB-Tk then the average protein similarity output by MAGpy was <95%. Genera were defined as novel if all MAGs which clustered at 60% AAI (49) were not assigned a genus by GTDB-Tk. Proposed names for new genera and species belonging to these genera were formulated based on the International Code of Nomenclature of Prokaryotes (50). To assess the abundance of our MAGs in other chicken populations, reads from Munk *et al.* (35) were downloaded from the European Nucleotide Archive (accession number: PRJEB22062), trimmed using cutadapt (51), aligned to the MAG database using BWA MEM and processed using Samtools.

Carbohydrate active enzymes (CAZymes) were identified by comparing MAG proteins to the CAZy database (52) using dbcan2 (version 7, 24^th^ August 2018). The abundance of CAZyme groups was then calculated as the sum of reads mapping to MAG proteins within each group after using DIAMOND (53) to align reads to the MAG proteins.

### Statistics and graphs

Univariate general linear models (GLMs) were performed in SPSS Statistics 21 (IBM) with gender, line and diet as fixed factors. All other statistical analyses were carried out in R (54) (version 3.5.1.). NMDS graphs were constructed using the Vegan package (55) and ggplot2 (56), using the Bray–Curtis dissimilarity. Boxplots were constructed using the ggplot2 package. UpSet graphs were constructed using the UpSetR package (57). Correlation coefficients, using R’s hclust function, were used to cluster samples and MAGs within heatmaps. PERMANOVA analyses were performed using the Adonis function from the Vegan package. The package DESeq2 (58) was used to calculate differences in abundance for individual MAGs, taxonomies and CAZymes. For MAGs, subsampling to the lowest sample coverage was performed prior to analysis by PERMANOVA and NMDS and before calculating the 1X and 10X coverage of MAGs in samples.

### Data availability

Paired-read fastq files have been submitted to the European Nucleotide Archive under project PRJEB33338. MAG fasta files have been submitted to Edinburgh DataShare (https://doi.org/10.7488/ds/2584).

## Results

### Assembly of 469 draft microbial genomes from chicken caeca

We produced 1.6T of Illumina data from 24 chicken samples, carried out a metagenomic assembly of single samples and also a co-assembly of all samples. 4524 metagenomic bins were created from the single-sample binning and 576 more were created from co-assembly binning. We were left with a total of 469 dereplicated genomes (99% ANI) with estimated completeness of ≥80% and estimated contamination ≤10% (**Supplementary figure 1**), 377 of which originated from the single-sample binning and 92 from the co-assembly. Of these, 349 had completeness >90% and contamination <5% (high-quality draft genomes as defined by Bower *et al.* (59)), 210 were >95% complete with <5% contamination and 47 MAGs were >97% complete with 0% contamination. The distribution of these MAGs (based on coverage) between the 24 samples can be found in **Dataset 1**. After dereplication to 95% ANI, 335 MAGs remained, representing species identified in our samples. Our dataset therefore contains 469 microbial strains from 335 species. 283 of these species and 460 of these strains were novel when compared to public databases **(Dataset 2)**.

**Dataset 2** contains the NCBI taxonomic assignment for each MAG along with the assembly characteristics and GTDB-Tk taxonomic assignments. **Dataset 3** contains comparative genomics information produced by MAGpy. **Figure 1** shows a phylogenetic tree of the MAGs. This was used to manually correct any errors in taxonomic identification. By far the most dominant phylum was *Firmicutes_A* (n=399), followed by *Firmicutes* (n=51), *Actinobacteriota* (n=10), *Proteobacteria* (n=3: all *Escherichia coli*), *Verrucomicrobiota* (n=2: genera *UBA11493* and *CAG-312*), *Bacteroidota* (n=1: *Alistipes sp. CHKCI003*), *Campylobacterota* (n=1: *Helicobacter_D pullorum*), *Cyanobacteriota* (n=1: order *Gastranaerophilales*) and *Desulfobacterota* (n=1: genus *Mailhella*). All members of *Firmicutes_A* belonged to the class *Clostridia*, which included the orders *Oscillospirales* (n=179), *Lachnospirales* (n=134), *4C28d-15* (n=42), *Christensenellales* (n=17), *TANB77* (n=10), *Peptostreptococcales* (n=9), *CAG-41* (n=5), *Clostridiales* (n=1), *UBA1212* (n=1) and one MAG which was undefined at order level (CMAG_333). All members of *Firmicutes* belonged to the class *Bacilli*; this included the orders *Lactobacillales* (n=21), *RF39* (n=20), *Erysipelotrichales* (n=8), *Exiguobacterales* (n=1) and *RFN20* (n=1). The *Actinobacteriota* were divided into two classes, *Actinobacteria* (n=5) and *Coriobacteriia* (n=5: containing only the order *Coriobacteriales*). The *Actinobacteria* class contained two orders: *Actinomycetales* (n=4) and *Corynebacteriales* (n=1). 97 MAGs were identified to species, 246 identified to genus, 115 identified to family, 10 identified to order and 1 identified to class. No MAGs were identified as Archaea.

**Figure 1:**
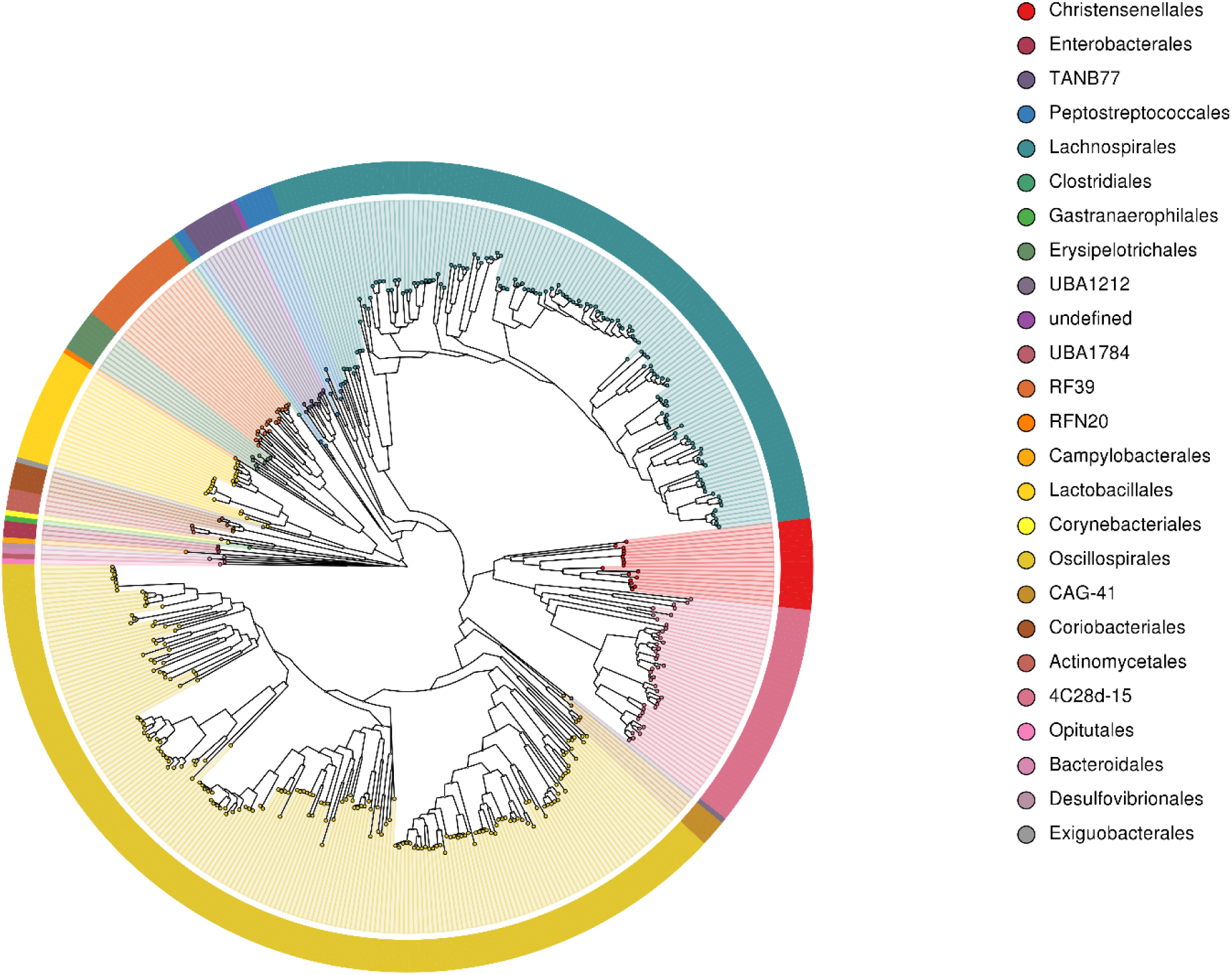
Phylogenetic tree of the 469 draft microbial genomes from the chicken caeca, labelled by taxonomic order, as defined by GTDB-Tk. Draft genomes labelled as “undefined” were only able to be assigned taxonomy at a higher level than order.

Of the MAGs that show greater than 95% ANI with an existing sequenced genome, several of these genomes have previously been identified in chickens. Our MAGs include 6 novel strains of *Anaeromassilibacillus sp. An250* (20), a novel strain of *Anaerotignum lactatifermentans* (60), a novel strain of *Blautia sp. An81* (20), 3 novel strains of *Drancourtella sp. An57* (20), a novel strain of *Enterococcus cecorum* (61), 2 novel strains of *E.coli* (14, 62, 63), 3 novel strains of *Eubacteriaceae bacterium CHKCI004* (64), a novel strain of *Eubacterium sp. An11* (20), two novel strains of *Faecalibacterium* spp. (20, 32), 7 novel strains of *Flavonifactor* spp. (20), 3 novel strains of *Gordonibacter* spp. (20), 1 novel strain of *Helicobacter pullorum* (65), 15 novel strains of *Lachnoclostridium* spp. (20), 6 novel strains of *Lachnospiraceae bacterium UBA1818* (32), 2 novel strains of *Massiliomicrobiota sp. An134* (20) and 5 novel strains of *Pseudoflavonifractor sp. An184* (20).

We also identified several Lactobacillli which have previously been isolated from the chicken gastrointestinal tract and have been suggested as potential probiotics in chickens, including 5 novel strains of *Lactobacillus crispatus* (66–68), 2 novel strains of *Lactobacillus gallinarum* (69), a novel strain of *Lactobacillus johnsonii* (70, 71), a novel strain of *Lactobacillus oris* (72), a novel strain of *Lactobacillus reuteri* (63, 66, 73) and a novel strain of *Lactobacillus salivarius* (63, 71, 74).

Our MAGs represent several putative novel species from 7 taxonomic classes: including 25 species of *Bacilli*, 252 species of *Clostridia*, 2 species of *Coriobacteriia*, 1 species of *Desulfovibrionia*, 1 species of *Lentisphaeria*, 1 species of *Vampirovibrionia* and 1 species of *Verrucomicrobiae*. These include 5 novel species of *Lactobacillus*. Our MAGs also contain 42 putative novel genera which contain 69 of our MAGs. We defined a genus as novel if all MAGs which clustered at 60% AAI were not assigned a genus by GTDB-Tk (**Dataset 4**). 40 of these novel genera belong to the class *Clostridia*, with over half belonging to the order *Oscillospirales*. One of the remaining novel genera contains one MAG which belongs to the *Bacilli* class (order *Exiguobacterales*) while the remaining genus belongs to the *Cyanobacteriota* (*Melainibacteria*), within the order *Gastranaerophilales*. Our proposed names for these genera and the species they contain can also be found in **Dataset 4**, alongside descriptions of their derivations. GTDB-Tk was unable to assign taxonomy to either of these genera at lower than order level, indicating that they may belong to novel bacterial families. It should also be noted that several genus-level MAG clusters do not contain any MAGs which were assigned a valid NCBI genus label but instead only received names defined by GTDB-Tk. For example, Group 16 (**Dataset 4**) is entirely constituted by MAGs of the genus *UBA7102*.

### Newly constructed MAGs are abundant in chicken populations across Europe

In order to assess the abundance of our MAGs in other chicken populations, we compared sequence reads generated from 179 chicken faecal, pooled, herd-level samples, collected from eight different countries across the European Union (35), to the 469 MAGs generated as part of this study. The read mapping rates can be seen in **figure 2**. Over 50% of the reads mapped to the MAGs in all samples; in seven out of eight countries the average read mapping rate was above 70%; and in Italy the average read mapping rate was above 60%.

**Figure 2:**
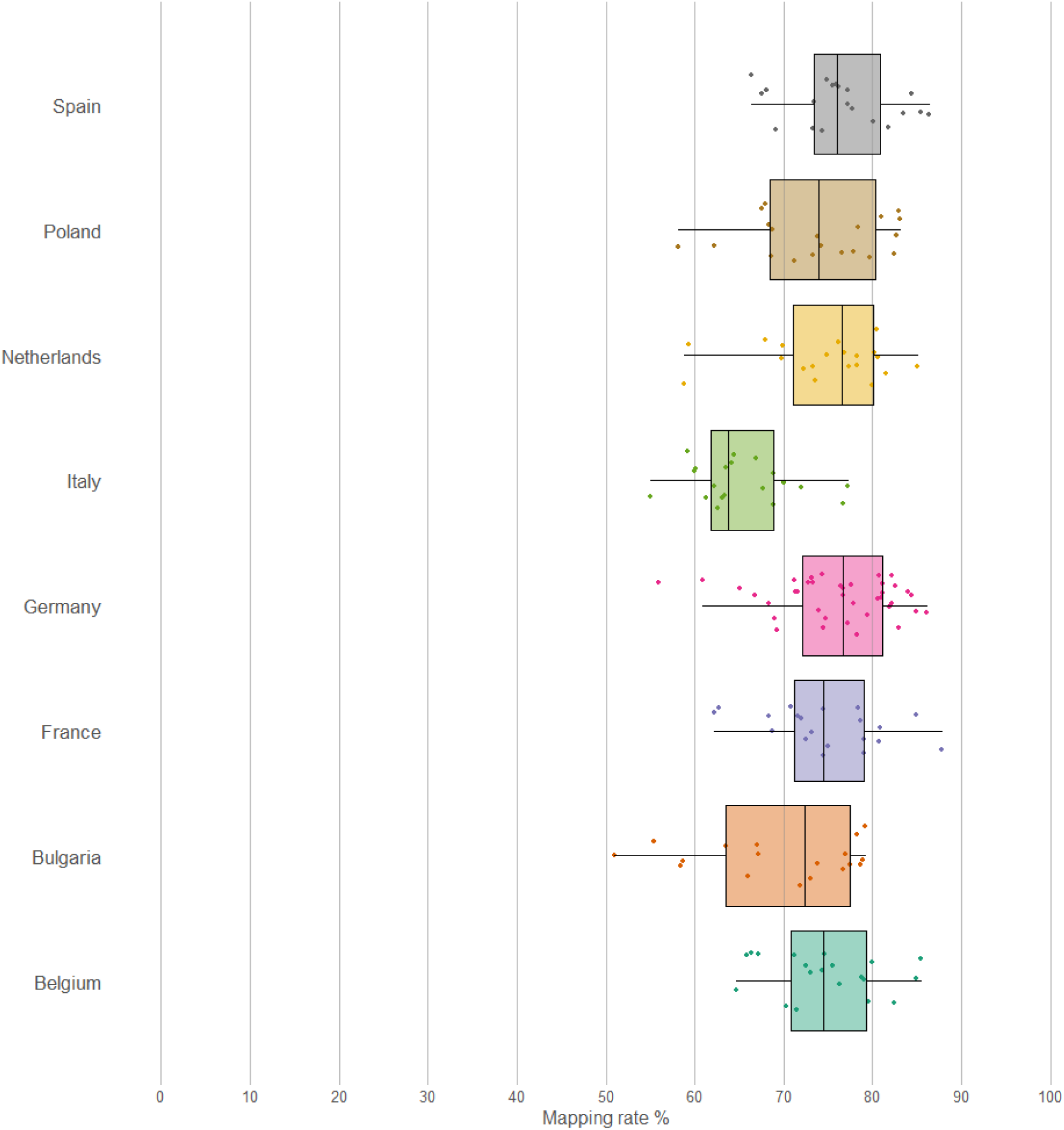
Read mapping rates of 179 chicken faecal samples, from 8 EU countries, against a database of the 469 MAGs

This demonstrates that our MAGs are representative of the chicken gut microbiome in populations throughout the EU, and account for the majority of reads in all cases. The abundance of the MAGs across the 179 samples can be seen in **figure 3**. Whilst there is clear structure in the data, samples do not appear to cluster by country, and the observed similarities may be explained by other factors not available, such as breed, age, or diet.

**Figure 3:**
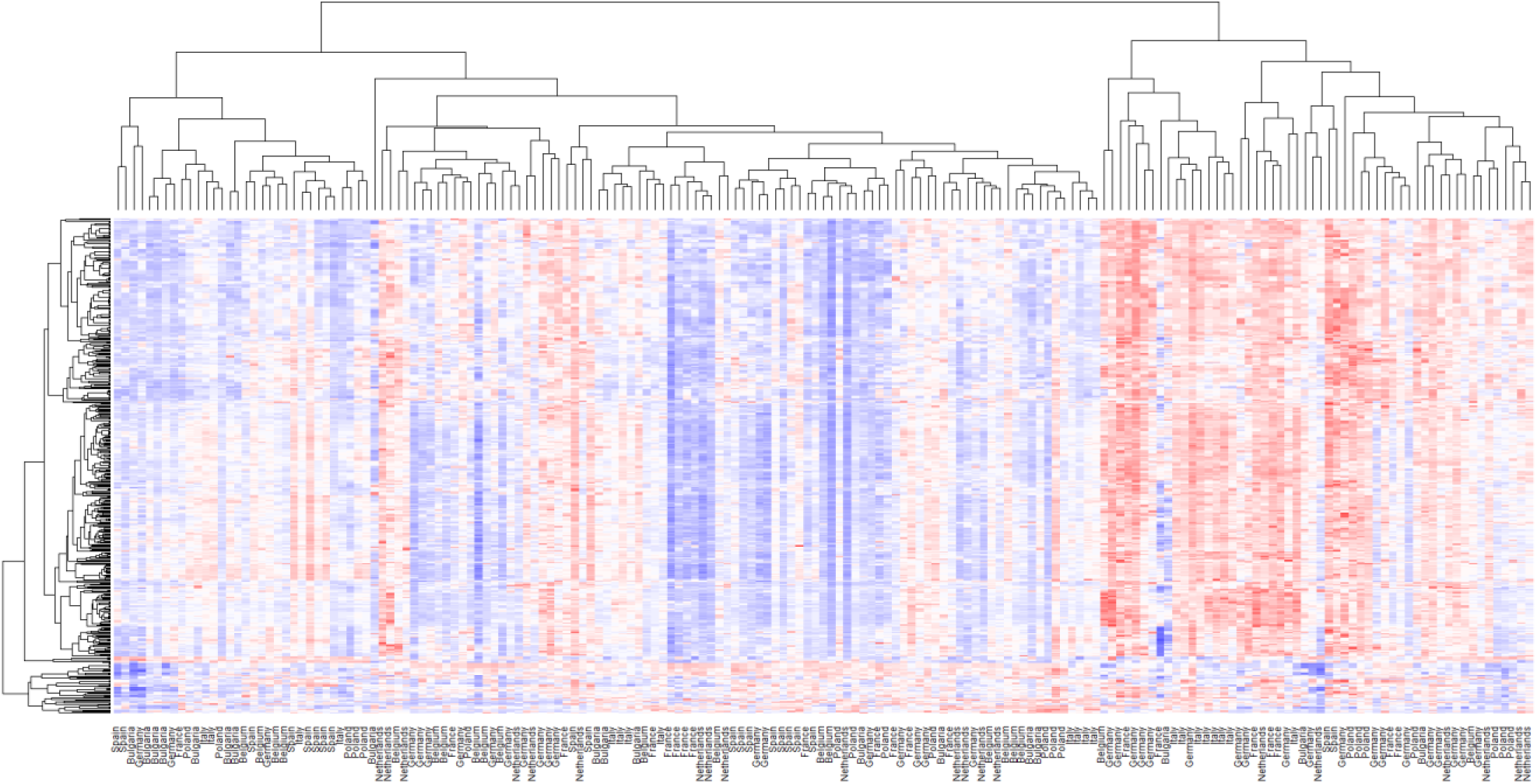
Abundance of 469 MAGs in 179 pooled chicken faecal samples from eight countries in the EU. Blue is low abundance, white medium, and red high abundance. Data are scaled within row.

### Presence of a core chicken caecal microbiota

125 MAGs were found to be present in at least 1X coverage in all of our samples and 4 of these MAGs were found to be ≥X10 in all of our samples: *Alistipes sp. CHKCI003* CMAG_6, uncultured *Bifidobacterium* sp. CMAG_55, uncultured *Bifidobacterium* sp. CMAG_59 and *Firmicutes bacterium CAG_94* CMAG_438. Only one MAG was found to be uniquely present in only one sample at ≥1X coverage: uncultured *Clostridia* sp. CMAG_391 in Chicken 16 (Ross 308: Vegetable diet). The distribution of MAGs between groups can be seen in **Figure 4.** 276 MAGs were on average present at at least 1X coverage in all groups and could therefore be described as a core microbiota shared amongst the chickens in our study.

**Figure 4:**
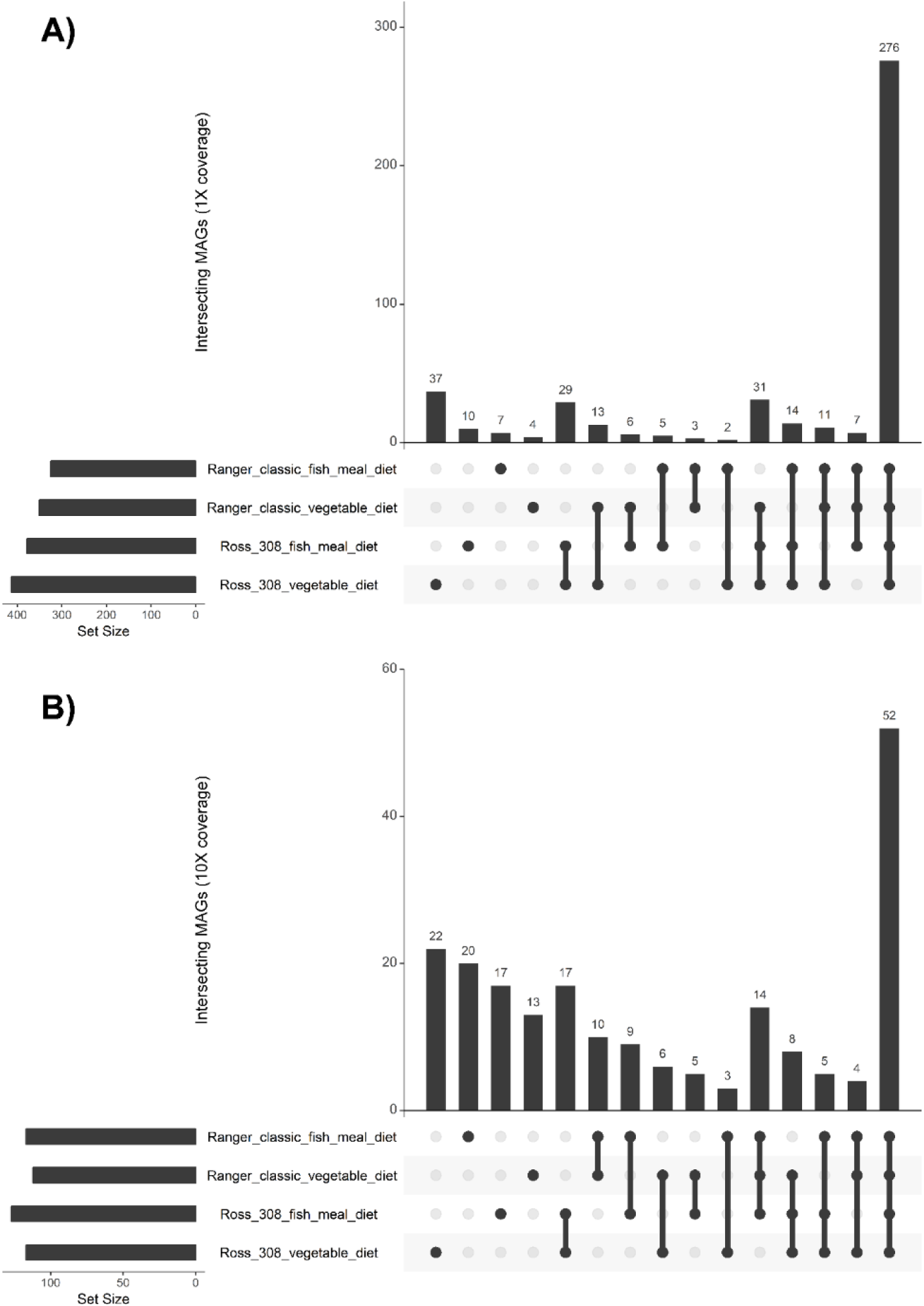
UpSet graphs showing the number of shared MAGs at A) average 1X coverage and B) average 10X coverage in the four chicken groups

### Differences in caecal MAGs based on chicken line and diet

When comparing samples based on the coverage of MAGs, significant clustering of samples by group can be observed when comparing all groups (PERMANOVA: P < 0.001); between chicken lines (All samples: PERMANOVA: P < 0.001; Within vegetable diet: PERMANOVA: P = 0.015, Within fish meal diet PERMANOVA: P = 0.0082)(**Figure 5**) and between diets (All samples: PERMANOVA: P = 0.008; Within Ross 308 line: PERMANOVA: P = 0.018; Within Ranger Classic line: PERMANOVA: P = 0.0043) (**Figure 5**). A significant interaction was also observed between line and diet (Line*Diet PERMANOVA: P = 0.038). Gender and DNA extraction batch were not found to have significantly affected the abundance of MAGs (PERMANOVA: P>0.05).

**Figure 5:**
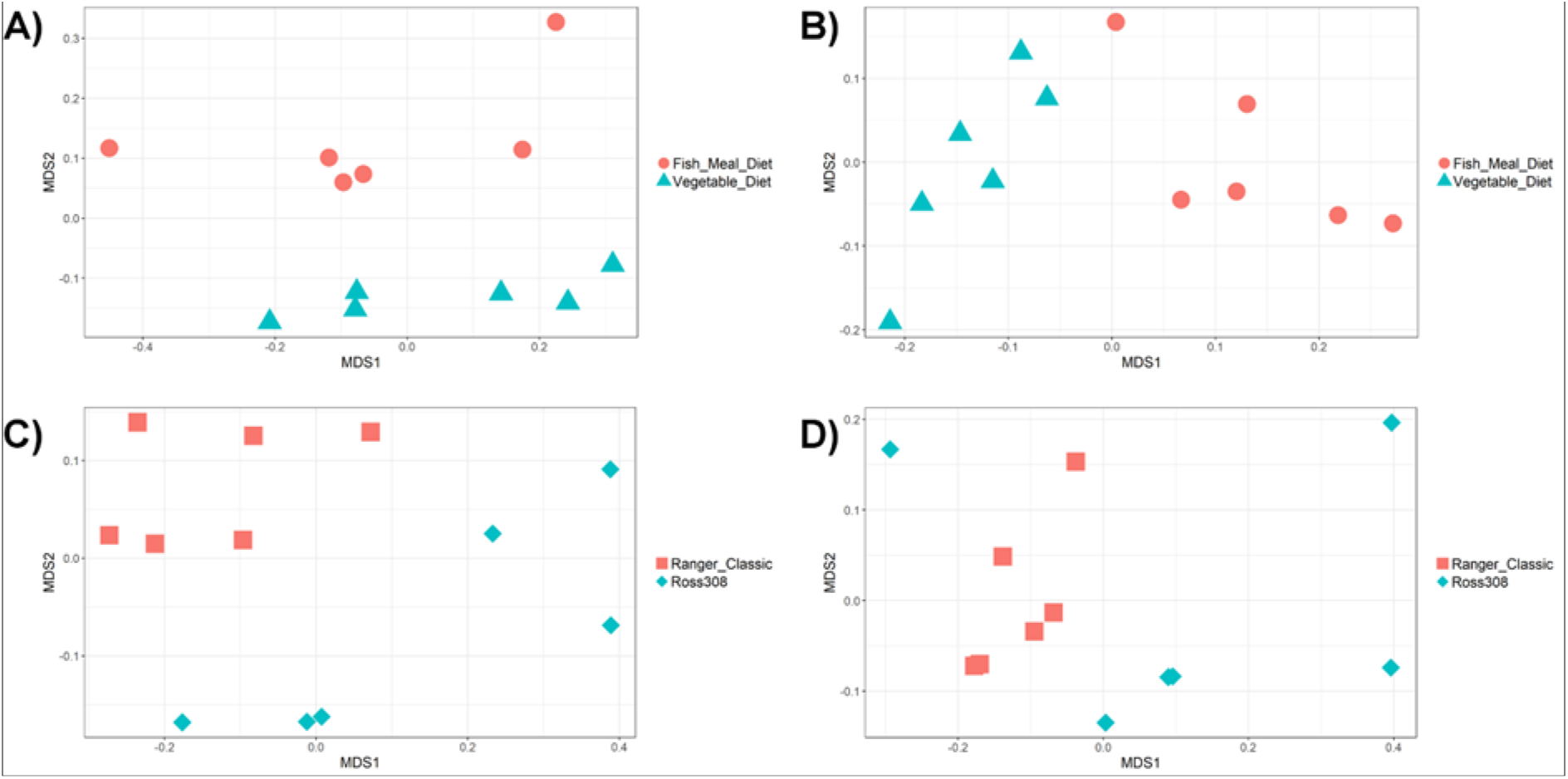
NMDS of chicken caecal samples clustered by proportion of MAGs (Bray-Curtis dissimilarity). A) Ross 308 birds clustered by diet (PERMANOVA: P = 0.018) B) Ranger Classic birds clustered by diet (PERMANOVA: P = 0.0043) C) Birds on a vegetable diet clustered by line (PERMANOVA: P = 0.015) D) Birds on a fish meal diet clustered by line (PERMANOVA: P = 0.0082).

MAGs which were significantly more abundant by coverage between groups were identified by DESeq2 (**Figure 6**); a full list of these MAGs can be found in **Dataset 5**. In Ross 308 birds, 43 MAGs were found to be differentially abundant between the two diets, while in Ranger Classic birds 45 MAGs were found to be differentially abundant. Several MAGs were found to be differentially abundant between the two lines when birds were consuming a vegetable diet (61 MAGs) or a fish meal diet (69 MAGs). 98 MAGs were found to be differentially abundant between lines when controlling for diet and 64 MAGs were found to be differentially abundant between diets when controlling for line.

**Figure 6:**
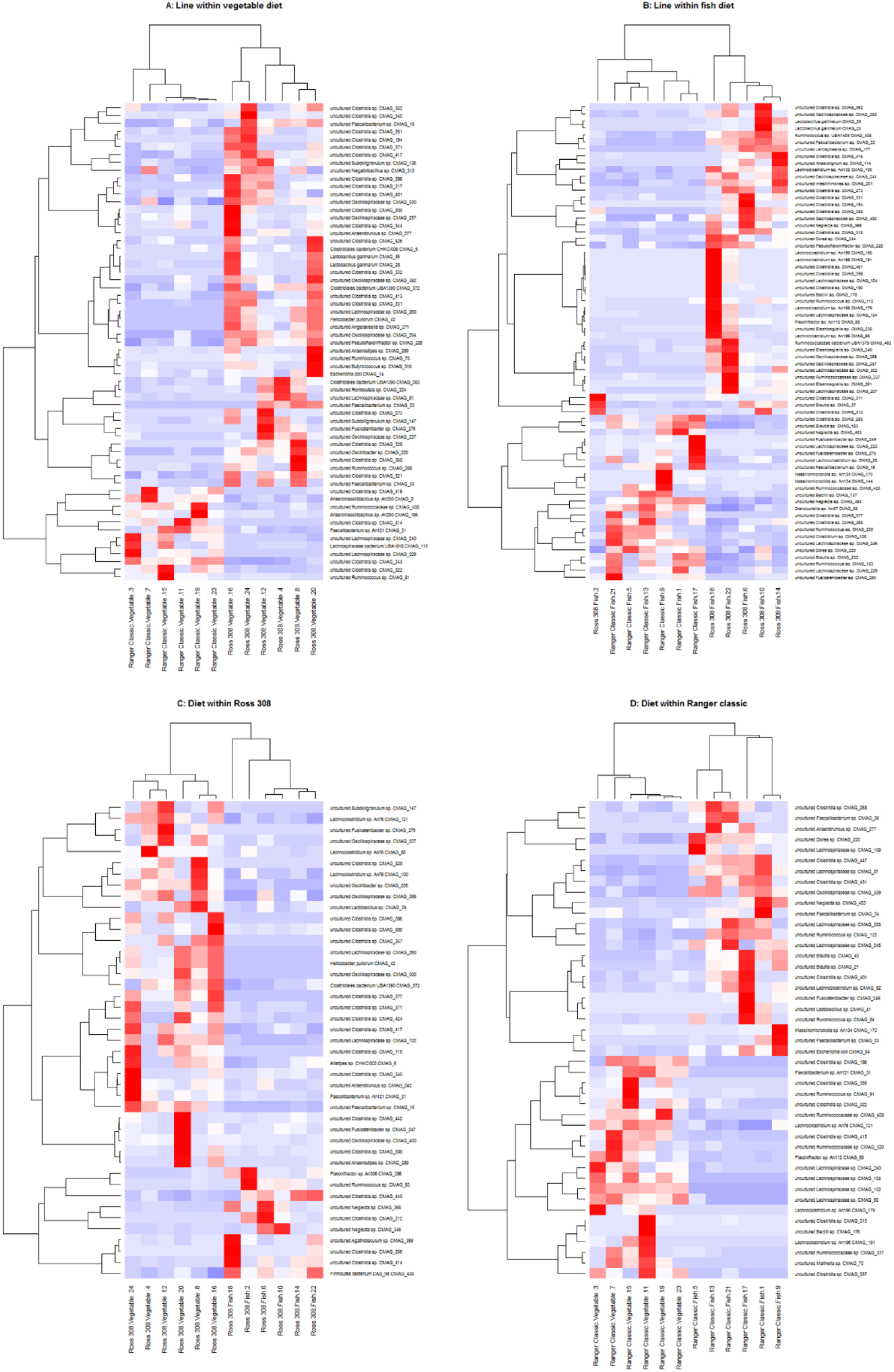
Heatmap showing the proportional coverage of MAGs which were significantly differently abundant between groups (Deseq2, P ≤ 0.05). Euclidean clustering was used to cluster MAGs and samples.

No MAGs were found to be significantly more abundant in both Ross 308 and Ranger Classic birds fed a fish meal diet, whilst four MAGs were found to be significantly more abundant in both Ross 308 and Ranger Classic birds fed a solely vegetable diet: uncultured *Lachnospiraceae* sp. CMAG_102, *Lachnoclostridium sp. An76* CMAG_121, *Faecalibacterium sp. An121* CMAG_31 and uncultured *Clostridia* sp. CMAG_357.

Eight MAGs were found to be significantly more abundant in Ross 308 chickens on both diets: uncultured *Pseudoflavonifractor* sp. CMAG_226, uncultured *Oscillospiraceae* sp. CMAG_257, uncultured *Clostridia* sp. CMAG_273 and uncultured *Clostridia* sp. CMAG_331, *Clostridia* sp. CMAG_194, *Lactobacillus gallinarum* CMAG_28, uncultured *Faecalibacterium* sp. CMAG_33 and *Lactobacillus gallinarum* CMAG_35. In contrast, only one MAG was found to be consistently more abundant in Ranger Classic birds on both diets (uncultured *Lachnospiraceae* sp. CMAG_229).

Lactobacilli are of particular interest to probiotic manufacturers. We found that both MAGs identified as *L.gallinarum* were more abundant in Ross 308 birds when controlling for diet, and four of the five MAGs identified as *L.crispatus* were more abundant in birds fed a diet with fish meal when controlling for chicken line.

One notable observation is the high amount of *Helicobacter pullorum* observed in the Ross 308: Vegetable diet group. While *H. pullorum* is often thought of as a pathogen, it has previously been isolated from the caeca of asymptomatic chickens (65) and carriage of *Helicobacter* by chickens is common in commercial flocks (75–77).

### Differences in CAZymes between lines and diets

Carbohydrate-active enzymes (CAZymes) are enzymes involved in the metabolism, synthesis and binding of carbohydrates. They are grouped by the CAZy database (52) into the following major groups: the auxiliary activities (AAs) class, carbohydrate-binding modules (CBMs), carbohydrate esterases (CEs), glycoside hydrolases (GHs), glycosyltransferases (GTs) and polysaccharide lyases (PLs). As their names suggest, CEs are responsible for the hydrolysis of carbohydrate esters while CBMs are responsible for binding carbohydrates. GHs and PLs are both responsible for cleaving glycosidic bonds, hydrolytically or non-hydrolytically respectively, while GTs are able to catalyse the formation of glycosidic bonds. The AA class are not themselves CAZymes but instead act in conjunction with them as redox enzymes. We compared the predicted proteins from our MAGs with the CAZy database using dbcan with the cut-offs E-value < 1e-18 and coverage > 0.35.

When clustering groups by the abundance of MAG derived CAZymes, all groups separate visually (**Figure 7**) but only the following differences were significant: Ross 308 birds were shown to cluster significantly by diet (PERMANOVA, P=0.021), and birds receiving a fish meal diet clustered significantly by line (PERMANOVA, P=0.0065). A significant interaction was observed between line and diet (Line*Diet PERMANOVA: P = 0.0051). Using DESeq2 we also found that the abundances of specific CAZymes differed between groups (**Figure 8**), full lists of which can be found in **Dataset 6**. We found several starch degrading enzymes to be differentially abundant between lines when controlling for diet, including GH13 subfamily 10, GH15, GH57, GH4 and GH31, and between diets when controlling for line, including GH13, GH13 subfamily 28 and GH13 subfamily 33. We also found that several CAZymes involved in metabolising cellulose and hemi-cellulose were differentially abundant between lines when controlling for diet, including GH5 (subfamilies 19, 37, 48, 44, 18), CE6, GH43 (subfamilies 30, 19, 29, 12), GH115, CE2 and GH67, and between diets when controlling for line, including GH5 (subfamilies 7 and 48) and GH43 (subfamilies 33, 4 and 35). Gender and DNA extraction batch were not found to have significantly affected the abundance of CAZymes (PERMANOVA: P>0.05).

**Figure 7:**
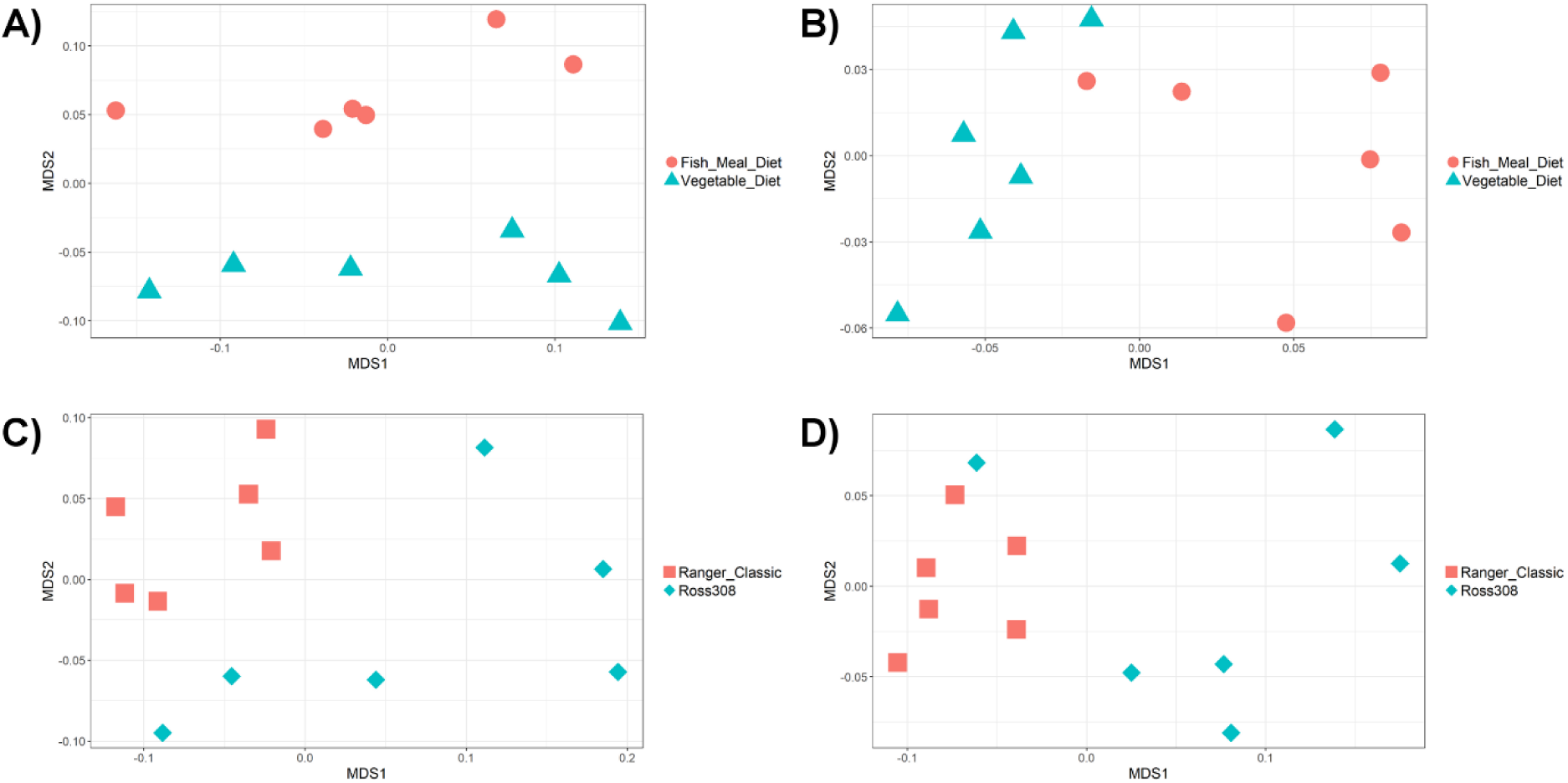
NMDS of chicken caecal samples clustered by abundance of MAG CAZymes (Bray-Curtis dissimilarity). A) Ross 308 birds clustered significantly by diet (PERMANOVA: P = 0.021) B) Ranger Classic birds did not cluster significantly by diet (PERMANOVA: P = 0.095) C) Birds on a vegetable diet did not cluster significantly by line (PERMANOVA: P = 0.061) D) Birds on a fish meal diet clustered significantly by line (PERMANOVA: P = 0.0065).

**Figure 8:**
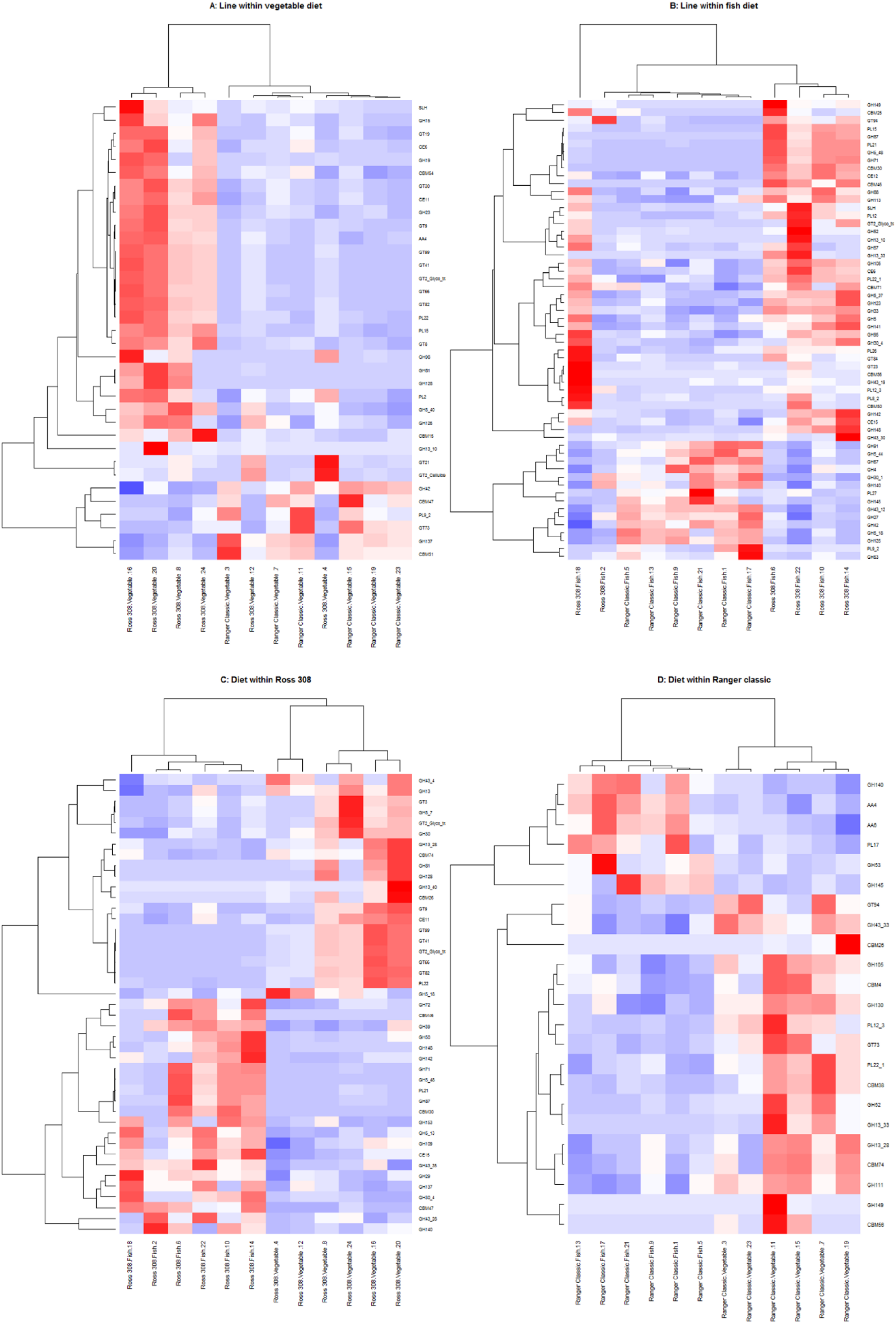
Heatmap showing the proportional coverage of MAGs which were significantly differently abundant between groups (Deseq2, P ≤ 0.05). Euclidean clustering was used to cluster MAGs and samples.

### Line and gender impact the weight of the chicken

As we did not monitor individual feed intake, we cannot comment on the feed-conversion ratio of these birds; however, when housed and fed as a group, there are clear statistical differences between the birds in terms of weight (**Supplementary figure 2**). Univariate GLMs with fixed factors of gender, line and diet were performed, with bird weight as the dependent variable. Both gender (P<0.001) and line (P<0.001) were found to significantly impact weight, as expected. Diet was not found to significantly affect bird weight overall (P=0.220). We did observe a significant increase in bird weight in Ranger Classic birds (P=0.007), of both genders, fed a fish meal diet; which was not observed in the Ross 308 birds (P=0.778).

## Discussion

It may be possible to increase chicken productivity by the manipulation of the chicken caecal microbiota. However, before this is possible we need to develop a good understanding of the types of bacteria present in the chicken and their nutritional function.

In this study we constructed 469 metagenome assembled genomes from chicken caecal contents, greatly expanding upon previous chicken caecal MAGs (25). 349 of our MAGs had completeness >90% and contamination <5% and can therefore be classed as high-quality draft genomes as defined by Bower *et al.* (59). Our MAGs include 460 novel strains and 283 novel species, including 5 novel *Lactobacillus* species. 97 MAGs were able to be identified to species level by GTDB-Tk and a further 246 could be identified to genus. We also identified 42 novel bacterial genera, 40 of which belonged to the class *Clostridia*. The remaining two genera belonged to the *Bacilli* class and the *Gastranaerophilales* order of *Cyanobacteriota*, and may also belong to novel taxonomic families. Our method of defining genera is conservative, as genera within different taxonomies may cluster at higher AAIs (49, 78, 79). We used GTDB-Tk instead of NCBI to assign taxonomies to our MAGs for the following reasons. The vast majority of our MAGs are members of the *Clostridia*, whose taxonomies are known to fit poorly with genomic data (80). Indeed, when we constructed a phylogenetic tree of our MAGs using NCBI classifications, we found many discrepancies between the taxonomic assignments and our tree (data not shown) resulting in the need for many manual corrections. However, using GTDB-Tk it was only necessary to manually correct one of our MAGs (CMAG_333) which was originally classified as a member of the Dehalobacteriia but clearly sat within the Clostridia in our tree. Our experiences reflect those of Coil *et al.* who found that the use of GTDB-Tk required less labour and reduced the need for subjective decisions in taxonomic assignment (81). The majority of our MAGs belonged to the orders *Oscillospirales* and *Lachnospirales*, members of the *Clostridia* class. The high abundance of *Clostridia* observed during our study correlates with several previous studies examining the chicken caecal microbiota (20, 82–87). This is likely the product of chicks being raised in an environment where they are not exposed to a maternal microbiota as feral hens and chicks exposed to an adult hen have microbiotas which are far less dominated by *Firmicutes* and contain higher abundances of *Bacteroidetes* (88, 89).

Within our dataset we found 276 microbes which were on average present at a minimum 1X coverage in all four of our groups, potentially indicating a core chicken microbiota. However caution must be taken as all of our chickens were raised in the same facility and samples were all taken at the same time-point, which will have limited the variability in microbes present. Chicken microbiota can vary across flocks (90), at different times in the bird’s life (91) and between free-range and intensively-reared chickens (92). To provide a truly representative dataset of chicken microbial genomes it would be necessary to sequence caecal samples from birds from multiple lines and raised under a variety of conditions. However, we do think it is likely that there is a core chicken caecal microbiota which is shared across sites and is irrespective of management conditions. Our comparison to chicken faeces samples from eight countries which were part of a pan-EU project on AMR demonstrates that our MAGs are abundant in chicken populations across Europe, and that these new genomes can account for the majority of reads in chicken gut microbiome studies. We also identified several novel *Lactobacillus* strains which have previously been posited as potential chicken probiotics, including *L.crispatus* (66–68), *L.gallinarum* (69), *L.johnsonii* (70, 71), *L.oris* (72), *L.reuteri* (63, 66, 73) and *L.salivarius* (63, 71, 74).

When analysing the abundance of MAGs between birds from different lines, consuming either a vegetable diet or a diet containing fish meal, we found significant differences in the microbial communities based on both line and diet. This agrees with previous studies where significant differences have been described in the intestinal microbiota of chickens from different lines, including those from faster and slower growing lines (93–95). Differences have also previously been observed in the microbiota when feeding chickens a diet supplemented with fish meal (33, 34). This correlates with differences observed in the weights of birds fed the fish meal diet. Ranger Classic birds fed a fish meal diet weighed significantly more than those fed a vegetable-only diet, whereas those was no significant difference between the weight of the Ross 308 birds fed on these two diets.

Examining those bacteria which were consistently significantly increased in a specific line regardless of diet or a specific diet regardless of line, the majority of these bacteria are novel species, therefore it is difficult to hypothesise why they are more abundant in particular bird lines or when birds are fed certain diets. Of those species that had previously been identified, the two *L.galinarum* strains were both consistently found to be more abundant in Ross 308 birds, while *Lachnoclostridium sp. An76* CMAG_121 and *Faecalibacterium sp. An121* CMAG_31 were found to be more abundant in birds on the vegetable diet. *L.gallinarum*, is a homofermentative and thermotolerant (69, 96) species which has previously been suggested as a potential chicken probiotic (67, 97, 98), while *Lachnoclostridium sp. An76* and *Faecalibacterium sp. An121* (20) have only very recently been discovered and are therefore not well characterised.

We are unsure why *H.pullorum* was observed in such high levels in the Ross 308: Vegetable diet group. We are unable to rule out contamination from the environment as our groups were housed in separate pens within the same room. We did not observe any negative health effects in this group, and the bacterium is very common in some flocks (65, 75–77, 99).

We wondered whether the differences in microbiota we observed between groups were associated with changes in the metabolic potential of the caecal microbial communities. Microbes isolated from the chicken caeca have previously been shown to have highly variable metabolic pathways (100, 101). We found that the abundances of certain MAG derived CAZymes involved in starch and cellulose degradation were significantly differently abundant between lines and diets. These molecules are highly abundant in the predominantly grain based diets fed to chicken. However, energy from starches and celluloses are not available to the chicken host unless these are first degraded into smaller carbohydrates by the gut microbiota, therefore differences between the ability of the caecal microbiota to degrade these molecules may lead to greater efficiency of energy extraction from feed (85).

It is also interesting to note that when analysing the abundance of MAG derived CAZymes in the chicken caeca, we only observed significantly separate clustering of birds by diet in the Ross 308 birds and by line in animals that were consuming the fish meal diet. This indicates that the differences in MAG abundances for these groups resulted in significantly different pools of metabolic genes. However, significant differences in MAG abundances were also observed for Ranger Classics on the two diets and for chickens of different lines consuming the vegetable diet, but this did not result in a significant difference in the total abundance of CAZymes. This finding serves to highlight that changes in microbiota community composition do not necessarily lead to significant changes in the total metabolic potential of that community, although it is possible more significant differences would be observed with a larger sample size. It is worth noting that while our Ross 308 vegetable diet group contained 4 males and 2 females and the other groups contained 3 males and 3 females, gender was found to have no impact on the abundance of CAZymes or MAGs and this therefore should not have impacted our results.

One outlier was observed in our data: Chicken 2 appeared to cluster separately by the abundance of its MAGs in comparison to other Ross 308 birds consuming a fish meal diet, supporting the idea that while diet and line are associated with differences in the microbiota, variation will still exist between birds of the same line consuming similar diets. It should also be noted that the individual feed intake of each bird was not measured, meaning that some birds may have consumed different quantities of food, which could lead to variation in their microbiota compositions.

In conclusion, through the construction of metagenome assembled genomes we have greatly increased the quantity of chicken derived microbial genomes present in public databases and our data can be used as a reference dataset in future metagenomic studies. While previous studies have demonstrated that *Clostridia* are very common in the chicken caeca, our study shows that within this class there is a wide diversity of species present, something which has perhaps been underestimated by culture based studies. To gain a mechanistic insight into the function of these bacteria and to capture the wide-diversity of bacteria present in chickens, large-scale culture based studies will be necessary.

## Acknowledgements

We would like to thank the staff at the Greenwood Building, Roslin Institute for the care of our animals. We would also like to thank Prof. Aharon Oren for aiding with the naming of new genera and species, and Denny Gorman for his help with sample preparations. The Roslin Institute forms part of the Royal (Dick) School of Veterinary Studies, University of Edinburgh. This project was supported by the Biotechnology and Biological Sciences Research Council, including institute strategic programme and national capability awards to The Roslin Institute (BBSRC: BB/P013759/1, BB/P013732/1, BB/J004235/1, BB/J004243/1). MJP is supported by the Quadram Institute Bioscience BBSRC-funded Strategic Program: Microbes in the Food Chain (Project No. BB/R012504/1) and its constituent project BBS/E/F/000PR10351 (Theme 3, Microbial Communities in the Food Chain) and by the Medical Research Council CLIMB grant (MR/L015080/1)

## Dataset legends

**Dataset 1:** Average coverage of MAGs in all samples. Coverage was calculated by mapping MAG scaffolds to the adaptor trimmed Illumina reads for each sample. The average coverage of the scaffolds from a MAG within a sample were taken as the average abundance of that MAG in the sample.

**Dataset 2:** Description of each chicken MAG (metagenome-assembled genome), including novelty of species or strain, NCBI_name, GTDB-Tk _taxonomy, CheckM completeness and contamination, assembly size (mb), N50, number of contigs, the longest contig length (bp) and the GC content.

**Dataset 3:** Taxonomy assigned by MAGpy to MAGs.

**Dataset 4:** Clustering of samples at 60% AAI to form genus clusters. Novel genera were defined as clusters of MAGs at 60% AAI which were not assigned a genus by GTDB-Tk

**Dataset 5:** MAGs which were identified as being significantly more abundant by DESeq2 between diets and lines.

**Dataset 6:** CAZymes which were identified as being significantly more abundant by DESeq2 between diets and lines.

