## Supplementary figures and tables for "Assembly of hundreds of novel bacterial genomes from the chicken caecum"

**Supplementary figure 1:**

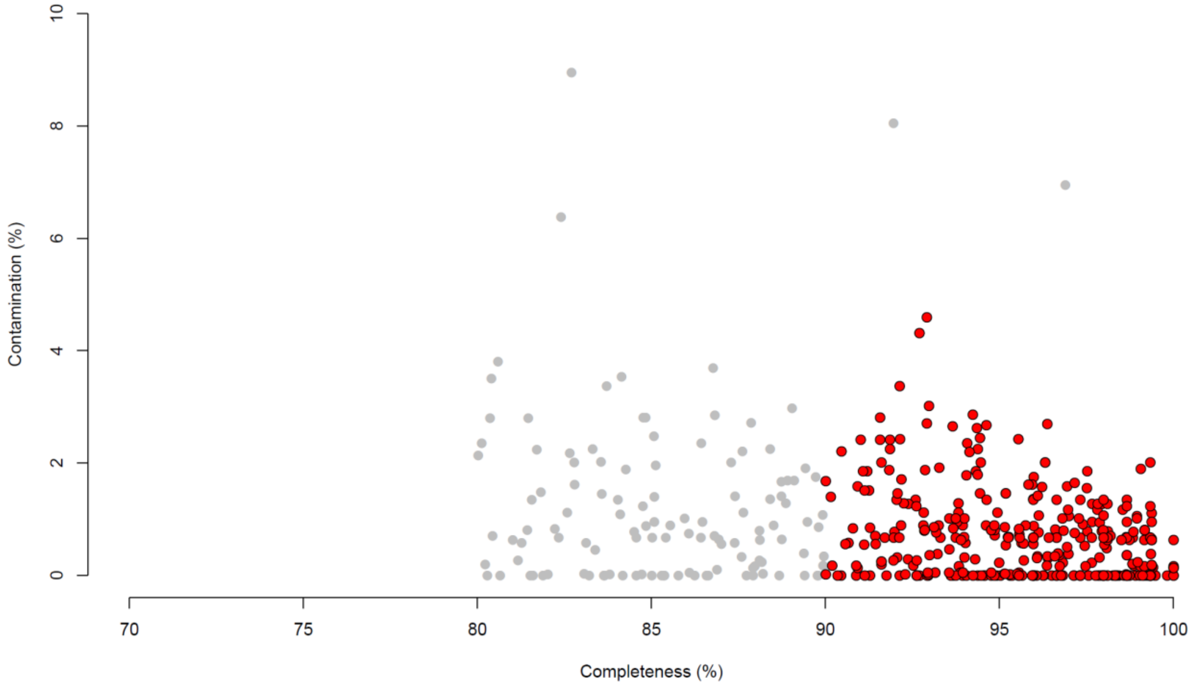


**Contamination and completeness of 469 dereplicated draft microbial genomes from the chicken caeca. Contamination and completeness were defined by the CheckM software. Red circles indicate genomes which are >90% complete with <5% contamination. Grey circles indicate genomes which are 80-90% complete with 5-10% contamination.**

**Supplementary figure 2:**

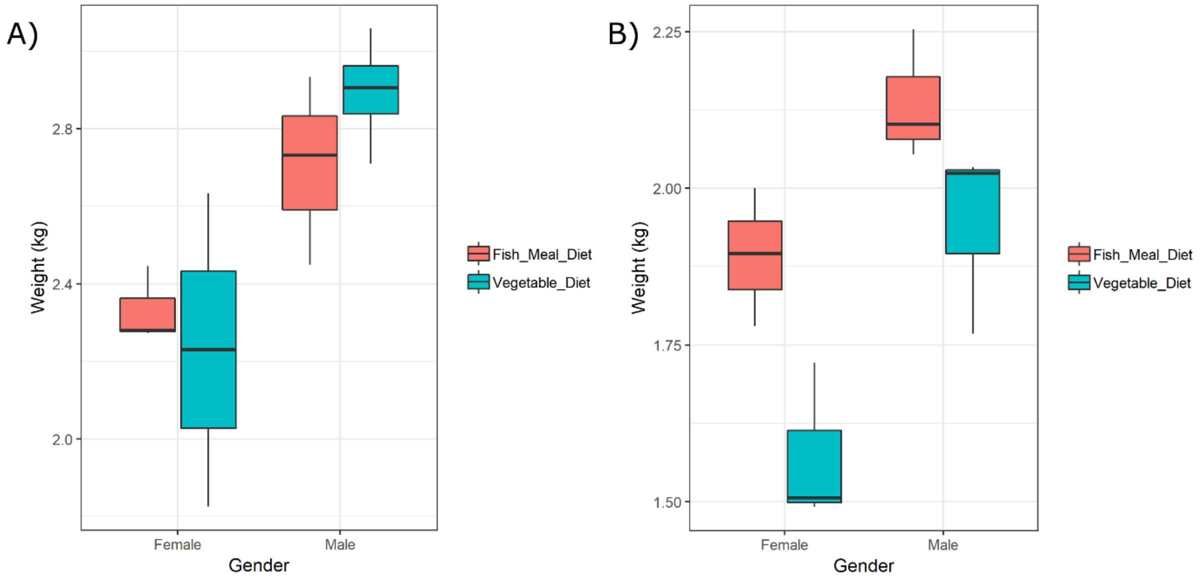


**Boxplots showing the weight of birds from either the Ross 308 (n=12) or Ranger Classic (n=12) line, fed either a vegetable diet or a diet which included fish meal.**

**Supplementary table 1: Chicken details**

| ID | Tag | Line | Diet | Gender | Body weight (kg) |
|---|---|---|---|---|---|
| Chicken_1 | 1472 | Ranger Classic | Fish | Female | 1.90 |
| Chicken_2 | 1677 | Ross 308 | Fish | Male | 2.73 |
| Chicken_3 | 1535 | Ranger Classic | Vegetable | Male | 2.03 |
| Chicken_4 | 1591 | Ross 308 | Vegetable | Male | 2.71 |
| Chicken_5 | 1445 | Ranger Classic | Fish | Female | 2.00 |
| Chicken_6 | 1724 | Ross 308 | Fish | Female | 2.28 |
| Chicken_7 | 1545 | Ranger Classic | Vegetable | Male | 2.02 |
| Chicken_8 | 1774 | Ross 308 | Vegetable | Male | 2.93 |
| Chicken_9 | 1531 | Ranger Classic | Fish | Male | 2.05 |
| Chicken_10 | 1738 | Ross 308 | Fish | Female | 2.27 |
| Chicken_11 | 1498 | Ranger Classic | Vegetable | Female | 1.49 |
| Chicken_12 | 1687 | Ross 308 | Vegetable | Female | 1.83 |
| Chicken_13 | 1423 | Ranger Classic | Fish | Female | 1.78 |
| Chicken_14 | 1675 | Ross 308 | Fish | Male | 2.45 |
| Chicken_15 | 1494 | Ranger Classic | Vegetable | Female | 1.72 |
| Chicken_16 | 1707 | Ross 308 | Vegetable | Male | 2.88 |
| Chicken_17 | 1473 | Ranger Classic | Fish | Male | 2.10 |
| Chicken_18 | 1764 | Ross 308 | Fish | Male | 2.93 |
| Chicken_19 | 1418 | Ranger Classic | Vegetable | Male | 1.768 |
| Chicken_20 | 1730 | Ross 308 | Vegetable | Male | 3.058 |
| Chicken_21 | 1421 | Ranger Classic | Fish | Male | 2.254 |
| Chicken_22 | 1640 | Ross 308 | Fish | Female | 2.446 |
| Chicken_23 | 1507 | Ranger Classic | Vegetable | Female | 1.506 |
| Chicken_24 | 1568 | Ross 308 | Vegetable | Female | 2.634 |

**Supplementary table 2: Formulation of fish meal and vegetable starter diets (Target Feeds Ltd, UK)**

| Raw materials | Broiler starter fish (%) | Broiler starter standard (vegetable) (%) |
|---|---|---|
| 1 Barley raw ground | 12.7 | 10.5 |
| 10 Wheat raw ground | 58.4 | 50.0 |
| 423 Soya ext hipro | 10.5 | 26 |
| 426 Full fat soya masham | 5 | 5 |
| 459 Provimi white fishmeal | 10 | 0 |
| 712 L-lysine Hcl | 0.35 | 0.4 |
| 713 DL methionine | 0.33 | 0.4 |
| 714 L-threonine | 0.15 | 0.15 |
| 716 L-Tryptophan | 0.02 | 0.00 |
| 810 Soya oil | 1.6 | 4 |
| 896 Limestone flour tru. 270 | 0.25 | 1.25 |
| 901 Monocalcium phosphate | 0.1 | 1.5 |
| 904 Salt | 0.05 | 0.25 |
| 906 Sodium bicarbonate | 0.15 | 0.15 |
| 3482 Br. Trials PMX. Starter 4K | 0.4 | 0.4 |

**Supplementary table 3: Formulation of fish meal and vegetable grower diets (Target Feeds Ltd, UK)**

| Raw materials | Broiler grower fish (%) | Broiler grower standard (vegetable) (%) |
|---|---|---|
| 1 Barley raw ground | 8.4 | 8.4 |
| 10 Wheat raw ground | 60.34 | 55.0 |
| 423 Soya ext hipro | 15.0 | 23.0 |
| 425 Full fat soya Cherwell | 5.0 | 5.0 |
| 459 Provimi white fishmeal | 5.0 | 0.0 |
| 712 L-lysine Hcl | 0.3 | 0.3 |
| 713 DL methionine | 0.31 | 0.35 |
| 714 L-threonine | 0.15 | 0.15 |
| 810 Soya oil | 3.25 | 4.5 |
| 900 Limestone Trucal 52 | 0.75 | 1.25 |
| 901 Monocalcium phosphate | 0.7 | 1.25 |
| 904 Salt | 0.25 | 0.25 |
| 906 Sodium bicarbonate | 0.15 | 0.15 |
| 3483 Br. Trials PMX. Gro/Fin 4K | 0.4 | 0.4 |

**Supplementary table 4: Nutritional info of fish meal and vegetable diets (Target Feeds Ltd, UK)**

| Nutrient | Broiler starter fish | Broiler starter standard (vegetable) | Broiler grower fish | Broiler grower standard (vegetable) |
|---|---|---|---|---|
| Oil EE | 4.5246 | 6.3685 | 5.8789 | 6.8528 |
| Protein | 21.7423 | 21.4436 | 20.2641 | 20.1993 |
| Fibre | 2.9826 | 3.1325 | 2.9279 | 3.0280 |
| Ash | 4.7791 | 6.0665 | 5.2399 | 5.6815 |
| Me-P | 12.7907 | 12.7703 | 13.0385 | 13.0441 |
| Tlysine | 1.4372 | 1.4343 | 1.2819 | 1.2682 |
| Avlysine | 1.3230 | 1.3356 | 1.1894 | 1.1829 |
| Meth | 0.6928 | 0.6900 | 0.6209 | 0.6260 |
| M+C | 0.9902 | 1.0155 | 0.9164 | 0.9373 |
| Threo | 0.9132 | 0.9013 | 0.8504 | 0.8497 |
| Trypt | 0.2509 | 0.2523 | 0.2235 | 0.2359 |
| Calcium | 0.9549 | 0.9805 | 0.9417 | 0.9304 |
| Phos | 0.6831 | 0.7289 | 0.6658 | 0.6575 |
| Av phos | 0.4908 | 0.4818 | 0.4598 | 0.4227 |
| Salt | 0.2991 | 0.3092 | 0.4024 | 0.3077 |
| Sodium | 0.1677 | 0.1749 | 0.2128 | 0.1763 |
| Vitamin A | 13.5000 | 13.5000 | 10.0000 | 10.0000 |
| Vitamin D3 | 5.0000 | 5.0000 | 5.0000 | 5.0000 |
| Vitamin E | 100.0000 | 100.0000 | 100.0000 | 100.0000 |

**Supplementary table 5: DNA extractions**

| ID | Sampling_date | DNA_extraction_date | DNA_extraction_batch |
|---|---|---|---|
| Chicken_4 | 27th June 2018 | 28th June 2018 | 1 |
| Chicken_8 | 27th June 2018 | 6th July 2018 | 2 |
| Chicken_12 | 27th June 2018 | 6th July 2018 | 2 |
| Chicken_16 | 27th June 2018 | 28th June 2018 | 3 |
| Chicken_20 | 27th June 2018 | 28th June 2018 | 3 |
| Chicken_24 | 27th June 2018 | 28th June 2018 | 1 |
| Chicken_2 | 27th June 2018 | 6th July 2018 | 2 |
| Chicken_6 | 27th June 2018 | 28th June 2018 | 3 |
| Chicken_10 | 27th June 2018 | 6th July 2018 | 2 |
| Chicken_14 | 27th June 2018 | 6th July 2018 | 2 |
| Chicken_18 | 27th June 2018 | 28th June 2018 | 3 |
| Chicken_22 | 27th June 2018 | 6th July 2018 | 2 |
| Chicken_3 | 27th June 2018 | 6th July 2018 | 2 |
| Chicken_7 | 27th June 2018 | 28th June 2018 | 1 |
| Chicken_11 | 27th June 2018 | 28th June 2018 | 1 |
| Chicken_15 | 27th June 2018 | 28th June 2018 | 3 |
| Chicken_19 | 27th June 2018 | 28th June 2018 | 3 |
| Chicken_23 | 27th June 2018 | 28th June 2018 | 1 |
| Chicken_1 | 27th June 2018 | 28th June 2018 | 1 |
| Chicken_5 | 27th June 2018 | 28th June 2018 | 3 |
| Chicken_9 | 27th June 2018 | 28th June 2018 | 1 |
| Chicken_13 | 27th June 2018 | 6th July 2018 | 2 |
| Chicken_17 | 27th June 2018 | 28th June 2018 | 1 |
| Chicken_21 | 27th June 2018 | 28th June 2018 | 3 |
